# Biomaterials direct functional B cell response in a material specific manner

**DOI:** 10.1101/2021.01.12.426347

**Authors:** Erika M. Moore, David R. Maestas, Chris C. Cherry, Jordan A. Garcia, Hannah Y. Comeau, Locke Davenport Huyer, Richard L. Blosser, Gedge D. Rosson, Jennifer H. Elisseeff

## Abstract

B cells are an adaptive immune target of biomaterials development in vaccine research but despite their role in wound healing have not been studied in tissue engineering and regenerative medicine. We evaluated the B cell response to biomaterial scaffold materials implanted in a muscle wound; a biological extracellular matrix (ECM) and synthetic polyester polycaprolactone. In the local muscle tissue, small numbers of B cells are recruited in response to tissue injury and biomaterial implantation. ECM materials induced plasmablasts in lymph nodes and antigen presentation in the spleen while the synthetic PCL implants delayed B cell migration and induced an antigen presenting phenotype. In muMt^−^ mice lacking B cells, the fibrotic response to the synthetic biomaterials decreased. Immunofluorescence confirmed antigen presenting B cells in fibrotic tissue surrounding silicone breast implants. In sum, the adaptive B cell immune response to biomaterial depends on composition and induces local, regional and systemic immunological changes.

## Introduction

The immune system is a central target of modern-day therapeutic strategies in cancer, autoimmunity and other diseases (*1, 2*). It has also emerged as an important contributor in regenerative medicine and wound healing (*3–6*). Biomaterial-based technologies are frequently a component of regenerative medicine strategies, where they serve as a scaffold for cell proliferation, differentiation and ultimately enable new tissue development (*7–9*). After implantation, biomaterials induce a foreign body response (FBR) (*10*). The immune response to biomedical implants is defined by a cascade of protein deposition, followed by neutrophil recruitment, macrophage response that results in fibrosis at the material interface (*10–12*). In the case of biomaterial scaffolds in regenerative medicine, the combined immune response to foreign biomaterial and localized tissue injury impacts the functional outcomes on the spectrum of regenerative repair to fibrosis. With technological development, the opportunities to better probe the diverse immune cell phenotypes that govern biomaterial-associated host response continue to grow.

B cells, a cell type of the adaptive immune system, are primarily recognized for their ability to produce antibodies (*13*). In effort to harness this effect, B cells have frequently been targeted with biomaterials, primarily in the form of micro- and nano-particles, in the context of vaccine development for infection response, treatment of autoimmune disease and cancer therapy (*8, 14, 15*). Other roles of B cells have received less attention in the context of the biomaterial response, including cytokine secretion, activation of other immune cells and antigen presentation. Although B cells make up a small percentage of cells types present local to biomaterial tissue implants, their potential importance in regenerative medicine is illustrated in B cell knockout models that resulted in reduced wound healing and reduced fibrosis (*11, 12, 16, 17*). Research on B cells thus far is mixed with divergent roles of B cells found in the biomaterial response and tissue repair. In one case, delivery of mature B cells promoted wound healing of skin incisions. In contrast, B cell presence correlated to fibrotic environments mediated through macrophage recruitment (*16, 17*). Further, reports of lymphoma development with synthetic breast implants also suggests B cell activity in detrimental fibrotic material responses (*18–21*). Considering that B cells traffic throughout the body, their response to biomaterials and wounds may hold important physiological implications on the systemic response to biomaterial implants.

In this work, we present an in-depth evaluation of the B cell response to biological and synthetic biomaterial scaffolds in a muscle injury. We characterize the kinetics and phenotype of B cells in response to a tissue extracellular matrix (ECM) biological scaffold and a synthetic polyester poly(caprolactone) (PCL) biomaterial implant. B cell phenotype changed in the regional lymph nodes and spleen depending on the material composition. Biological scaffolds induced an earlier B cell infiltration to the muscle wound environment, germinal center formation, and plasmablast development in the draining lymph node. In contrast, PCL materials showed delayed B cell infiltration and primarily induced a phenotype characterized by increased antigen presentation. Lack of B cells in a muMt^−^ KO model reduced the fibrotic response to PCL and decreased expression of inflammatory genes associated with fibrosis.

## Results

### Biomaterial scaffolds alter the B cell response to injury in the local tissue and regional lymph nodes

To characterize the B cell response to biomaterials, we performed a volumetric muscle loss injury on C57BL/6 wildtype (WT) mice and implanted two materials, submucosal-intestine extracellular matrix (SIS-ECM) or PCL. SIS-ECM promotes a pro-healing type 2 immune response characterized by interleukin-4 (IL-4) secretion (*5, 22*), while PCL promotes a type 17 immune response characterized by IL-17 secretion that yields the generation of senescent cells and a fibrotic capsule (*23*). To assess response phenotype, we performed multiparametric flow cytometry on the injured muscle tissue with and without biomaterials and the corresponding draining lymph node in comparison to naïve tissue. The number of CD45^+^ immune cells that migrated into the wound increased with implantation of biomaterial as expected based on previous studies (Fig. S1A). To assess B cells in the implants and tissue, we evaluated B220 (CD45R) and CD19 to differentiate mature and resting B cells (*24, 25*). B220 is a pan B-cell marker in mice while CD19 expression correlates to B cell developmental stage (*24, 25*). B cells expressing CD19^+^ B220^−^, CD19^−^B220^+^, and CD19^+^B220^+^ comprised a small percentage (3-12%) of the CD45^+^ cells in the muscles containing biomaterials and evolved over time (Fig. S1B).

The kinetics of CD19^−^B220^+^ B cell infiltration into the muscle injury depended on the presence and type of biomaterial (Fig. 1A). At 3-days post-surgery, the earliest time point analyzed, SIS-ECM recruited the greatest number of CD19^−^B220^+^ B cells to the muscle tissue (21570 ± 7452), compared to significantly lower numbers in the saline control (9064 ± 3077). In the SIS-ECM environment, B cells continue to infiltrate the injury environment over time, with a peak in cell number between 5 days and 1 week after injury. In contrast, PCL treated wounds reached peak B cell infiltration at 1 and 3-weeks after injury. At week-3 post-injury, PCL implantation recruited more B cells (11355 ± 1113) when compared to SIS-ECM (6501±1691) and saline (4682 ± 1819) treated groups. At 6-weeks post-injury, B cell numbers in the tissue were low and was similar in all treatment groups.

**Fig 1.**
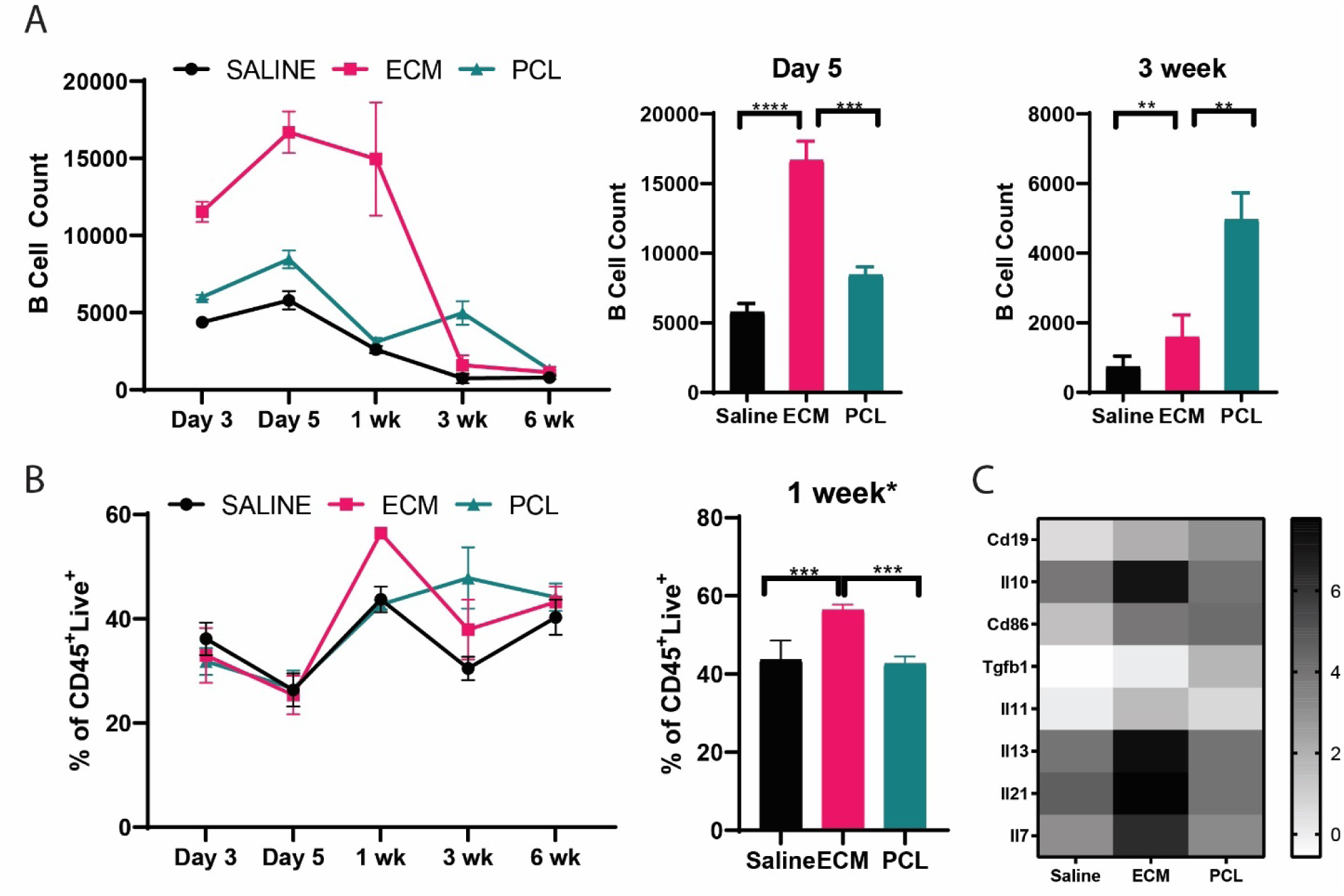
Biomaterial implants have diverging influences on B cells in quad tissue and in lymph node. (**A**) Flow cytometry counts of a subset of B cells (defined as CD19^−^B220^+^) in quad over time post injury. ECM recruits a peak number of B cells at Day 5 and PCL peaks at both 1 week and 3 weeks post-injury. (**B**) Flow cytometry assessment of B cells in the lymph node (represented as a percentage of CD45^+^Live^+^ cells) following injury and biomaterial implant. (**C**) Heatmap of Nanostring gene expression of B cell effector functions at 1 week after injury in the lymph node. Data are mean ± SD, n=4, Two-way analysis of variance (ANOVA) with subsequent multiple comparison testing. ANOVA [A, B: ****p<0.0001, ***p<0.001, **p<0.01].

B cell changes in the draining inguinal lymph node (ILN) reflected observed changes in the muscle tissue over time (Fig. 1B). Injury alone increased the total numbers of CD19^+^B220^+^ B cells in the ILN (Fig. S1C). The percentage of CD19^+^B220^+^ B cells in the ILN significantly increased at 1-week post-injury in muscle tissue treated with ECM compared to saline and PCL treatment (Fig. 1B). PCL treatment resulted in the greatest percentage CD19^+^B220^+^ B cells in the ILN at 3-weeks post injury. By 6-weeks, the percentage of B cells in the ILN was similar in all groups.

We further evaluated B cells using multiplex gene expression analysis of B cells sorted from the ILN 1 week after injury. ECM-treated B cells differentially upregulated B-cell phenotypic genes compared to injury alone (Fig. 1C, full expression data in data file S1). ECM and PCL treatment also upregulated genes associated with B and T cell interactions, including *Ctla4, CD86*, and *CD79a* when compared to injury controls (Fig. S2A). ECM treatment upregulated expression of Prdm1 (data file S1) that encodes the Blimp-1 gene, an regulator plasma cell formation, immunoglobulin secretion, and generation of long-lived plasma cells (*26*). PCL treatment induced a downregulation of Prdm1 expression, suggesting that ECM treatment supported more generation of plasma cells. Both biomaterial implant conditions upregulated antigen processing-related genes such as *CD1d1, Relb1, Icam1*, and *H2-DMb1* in B cells (Fig. S2B). However, ECM treatment resulted in reduced expression of genes associated with somatic hypermutations such as *fas* and *tirap*, while PCL downregulated *il7* (data file S1), a gene associated with B cell proliferation and survival (*27*).

### Single cell RNASeq of B cells reveals antigen specific responses and plasma cell generation after ECM treatment

To further characterize the B cell response to biomaterial scaffolds, we performed 10X 5’ single cell RNA Sequencing with transcriptome and B cell receptor sequencing on B cells isolated from the inguinal lymph nodes 3 weeks after biomaterial implantation. Visualization of cells using the UMAP (uniform manifold approximation and projection) dimensional reduction algorithm differentiated cell populations based on B cell identifiers, *CD19* and *Ms4a1* (*28, 29*) (Fig. S3A). Unbiased clustering algorithms categorized B cells into four clusters (Fig. 2A); 2 clusters in which the cells were largely undifferentiated (clusters 0,1), one cluster containing B cells with a type 1 interferon phenotype (cluster 2) and one cluster of B cells characterized by plasma cell formation (cluster 3) (Fig. 2A). ECM treatment upregulated plasma cell associated genes found in cluster 3 (Fig. 2A) including *Ighg1, Jchain*, and *Aicida*, all genes associated with germinal center formation, plasma cell generation, and class switching of B cells.

**Fig 2.**
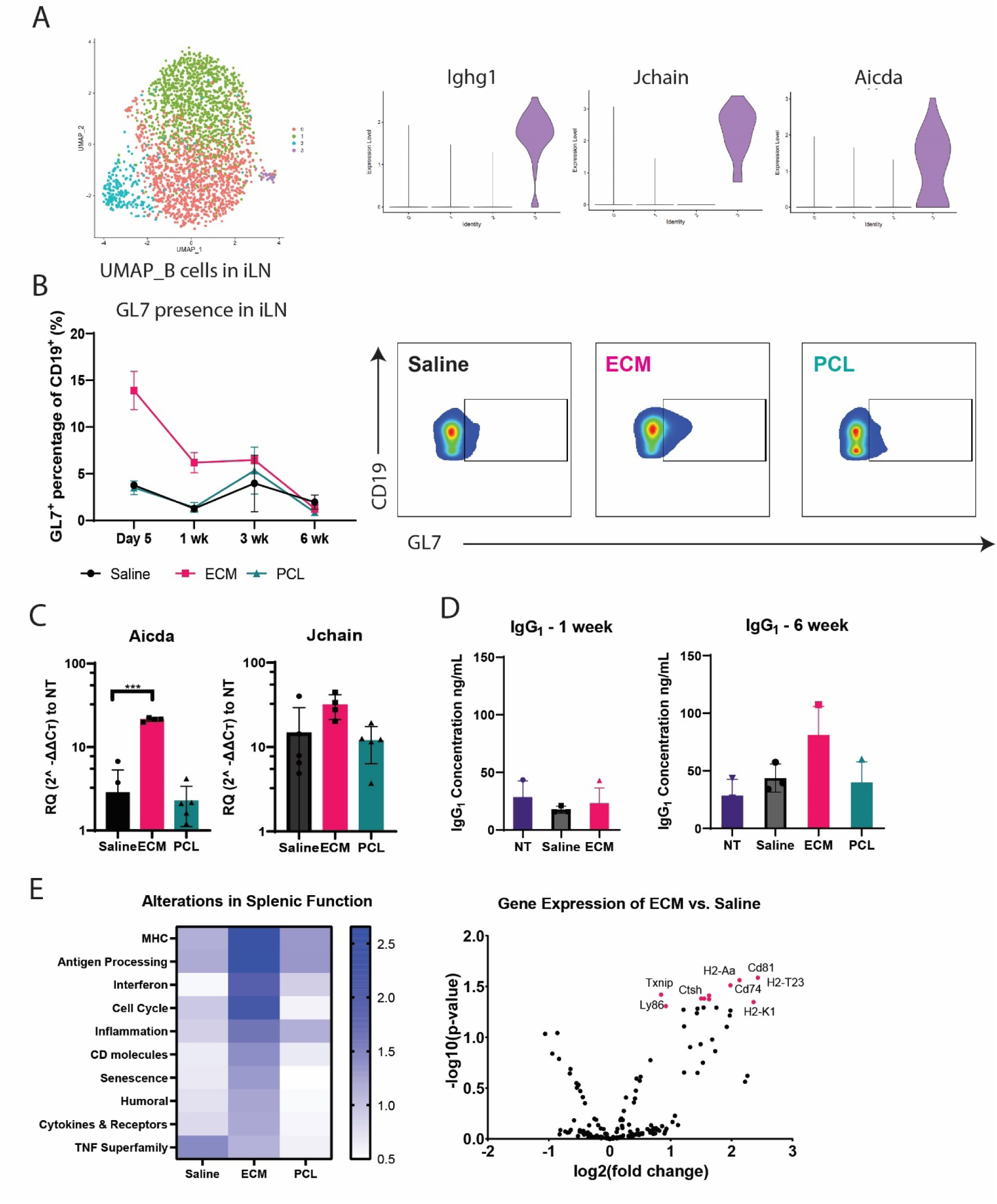
ECM induces plasmablasts and germinal center formation. (**A**) Dimensional reduction projection of B cells onto two dimensions using UMAP. Cells are colored by cluster. Violin plots of cluster gene expression of surface markers identified by differential expression analysis demonstrating genes highly expressed in cluster 3 associated with plasmablast generation. (**B**) Flow cytometry counts of GL7^+^ B cells (defined as GL7^+^CD19^+^) in the draining lymph node following injury +/- biomaterial. Flow cytometry plots of the Day 5 GL7 on B cells comparing Saline, ECM and PCL. (**C**) Gene expression using qRT-PCR at 1-week expression of *aicda* and *Jchain*. (**D**) Serum analysis of IgG1 at 1 week and 6 weeks post-injury. (**E**) Heatmap and volcano plot of Nanostring gene expression of B cell altered pathway functions at 3 weeks after injury in the spleen. Data are mean ± SD, n=4, Two-way ANOVA with subsequent multiple comparison testing [C: ***p<0.001].

To validate the plasma cell-like population specific to the ECM condition found in the single cell analysis, we probed expression of *Aicda* and *Jchain* in the lymph node 1-week following injury with quantitative PCR and assessed GL7, a common marker for germinal center B cells (Fig.2B,C). *Jchain* is expressed on all plasma cells and *Aicda* is expressed by B cells during germinal center development (*30, 31*). ECM induced a 10-fold increase in *Aicda* expression compared to saline and PCL treatment groups (Fig. 2C), and a similar induction of increased expression of *Jchain* expression.

To further probe the single cell cluster 3, we conducted multicolor flow cytometry, gene expression analysis, and immunoglobulin serum assessment. GL7, germinal center B cell marker, increased in the inguinal lymph node in response to ECM biomaterial implantation. ECM stimulated the greatest increase in GL7^+^ cells as a percentage of the total CD19^+^ population at 5 days after injury (Fig. 2B); GL7^+^ cells comprised 13.9 ± 2.0% of cells present in the ILN after treatment with ECM, compared to 3.5 ± 0.7 and 3.7 ± 0.2% in injuries treated with PCL and saline, respectively. Immunoglobulin serum analysis assessment further suggested differences in antibody production based on biomaterial treatment. ECM induced elevated levels of IgG1 after 6-weeks, indicating class switching had occurred in ECM treated groups compared to saline and non-injured (NT) groups (Fig. 2D). IgM increased in all groups 6-weeks after injury (Fig. S3B).

Antigen-specific B cell activation occurs through the B cell receptors (BCR) so the presence of BCR clones provides insight into potential antigen specificity of the B cell response to biomaterials. V(D)J library processing was conducted to investigate changes in immunoglobulin heavy-chain VH recombination and repertoires in the single cell B cell evaluation. ECM treatment produced the most clones compared to other groups (Fig. S4), suggesting ECM stimulates B cells to undergo somatic hypermutation (SHM) to create a diverse repertoire of clones. As ECM is a mixture of xenogeneic polysaccharides and proteins, the development of ECM specific clones is expected (*32*). However, clonal development was also observed in the PCL-treated group and in saline-treated control animals to a less marked degree.

Since B cells in the spleen can provide information on the systemic response (*33*), we sorted B cells from the spleen and performed multiplex gene expression analysis 3-weeks after biomaterial implantation. There was a significant difference in the gene expression profile of splenic B cells depending on biomaterial treatment. Splenic B cells from mice implanted with the ECM biomaterials upregulated genes associated with antigen processing, major histocompatibility complex (MHC) presentation, and pathways associated with cell cycle genes when compared to saline and PCL groups (Fig. 2E). MHC genes upregulated in ECM treatment group include *H2-K1, H2-Aa* and *H2-T23* compared to B cells sorted from saline-treated injuries. Finally, ECM treatment corresponded with upregulated CD molecule presentation, humoral associated genes and cytokines/receptors (Fig. 2E). In contrast, splenic B cells from mice treated with PCL significantly upregulated *S100a8*, an alarmin typically associated with environmental inflammation (*34, 35*) compared to saline treatment (Fig. S5).

### B cells increase antigen presentation in response to PCL and contribute to fibrosis

PCL implantation resulted in two peaks of B cell infiltration into injured muscle tissue: day-5 and 3-weeks post-implantation. The second B cell infiltration at 3-weeks post-implantation occurred as the other treatment groups resolved inflammation and B cell numbers decreased (Fig. 1A). B cell expression of B220^+^and CD19^+^, markers of B cell maturity that contribute to wound healing increased in the tissue with implants compared to injury alone at 3 weeks (*16*) (Fig. 3A). PCL treatment significantly increased the number of B220^+^ B cells in the tissue, in contrast to ECM and saline-treated animals (Fig. 3B). Furthermore, PCL implantation resulted in significantly higher numbers of antigen presenting MHCII^+^ B220^+^ cells in the tissue 3-weeks after implantation compared to the other conditions (Fig. 3B). PCL treatment further correlated with plasma B cell phenotypes (CD138^+^), seen with increased recruitment of CD138^+^B220^+^ B cells compared to ECM (Fig. S6A).

**Fig 3.**
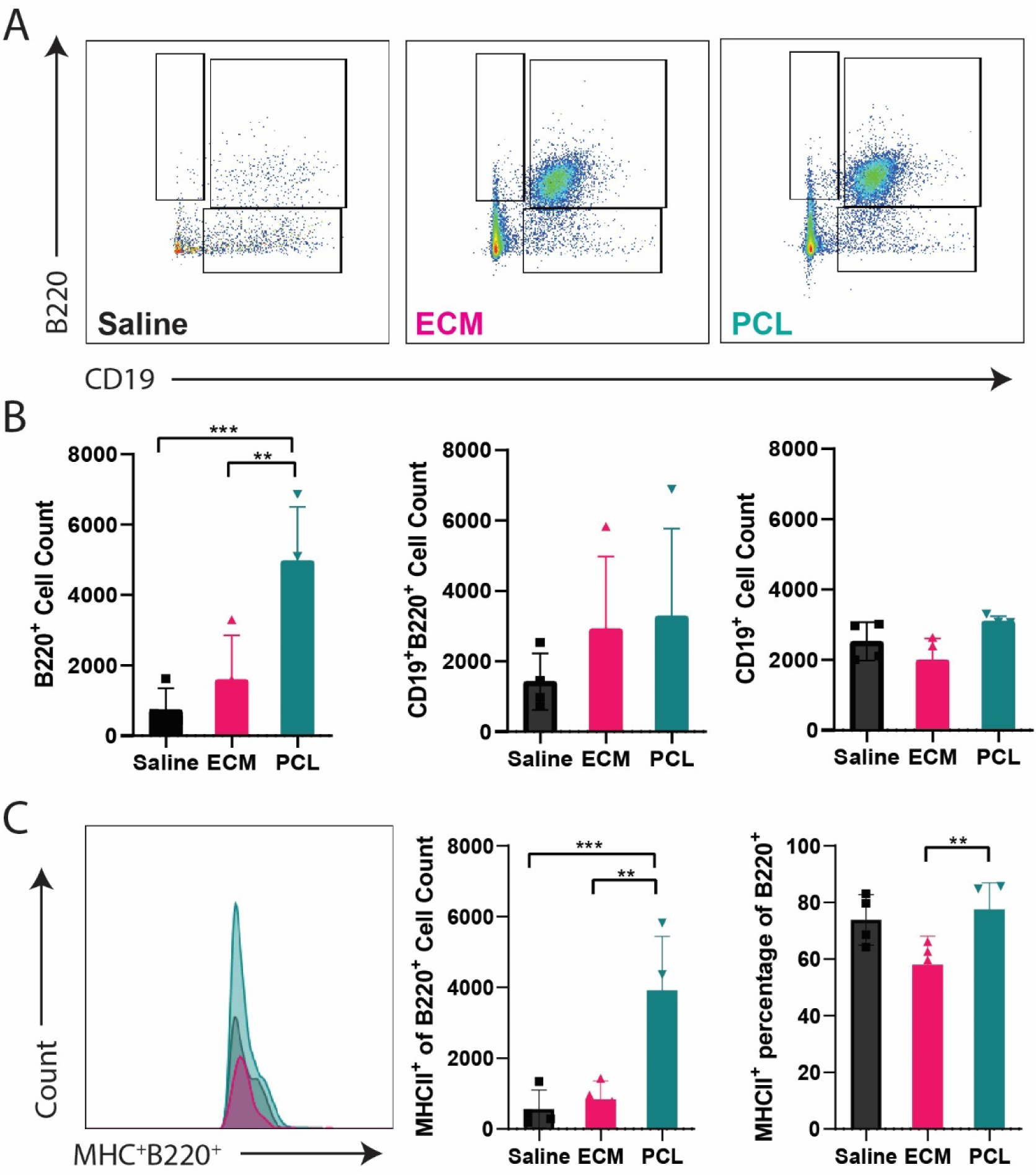
PCL induces B cell antigen presentation in the quad tissue. (**A**) Flow cytometry of B cell gating as a function of CD19 vs. B220 in the quad tissue after injury. (**B**) Flow cytometry counts of B cell subsets: CD19^+^B220^−^, CD19^−^B220^+^, and CD19^+^B220^+^ in the quad tissue 3 weeks post injury. (**C**) MHCII^+^B220^+^ cell population in quad tissue at 3 weeks post-injury. Data are mean ± SD, n=4, Twoway ANOVA with subsequent multiple comparison testing [B,C***p<0.001, **p<0.01].

To evaluate the functional impact of B cells in the FBR, we implanted the biomaterials in muMt^−^ mice, a homozygous mutant mouse strain that lacks mature B cells. In the muMt^−^ mice, there were no differences in numbers of CD4^+^ or CD8^+^ cells compared to WT at 1-week post-injury with biomaterials ECM or PCL (Fig. S7A). NK1.1 and γδ T cells decreased in the ECM biomaterial response at 1-week post-injury when compared to the WT (Fig. S7B). At 6-weeks post-injury, there was no statistical difference in cytokines associated with the PCL pro-fibrotic response (IL-17) in the muMt^−^ mice (Fig. S7C). Histological assessment of fibrosis around the PCL implants confirmed the reduced fibrosis in the MuMt^−^ mice. PCL in MuMt^−^ tissue injury yielded reduced collagen matrix deposition via Masson’s Trichrome compared to WT with PCL (Fig 4A). Additionally, there were significant differences in genes associated with fibrosis and the FBR. In the MuMt^−^ mice implanted with PCL, *S100a4, col1a1, col3a1* and *TGFβ* expression significantly decreased compared to WT mice implanted with PCL (Fig. 4B). *Il23* and *p21* also significantly decreased in the MuMt^−^ mice (Fig. 4B).

**Fig 4.**
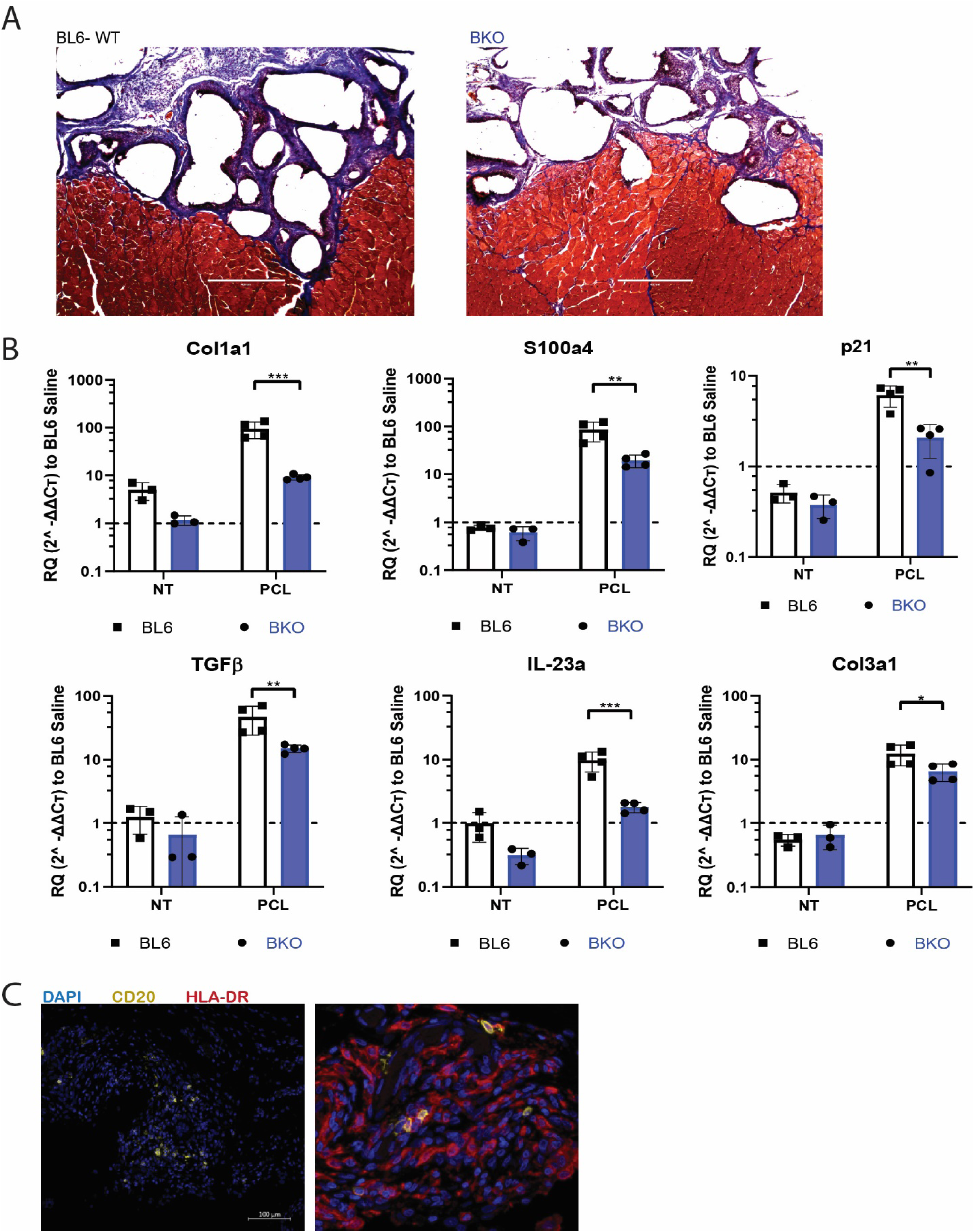
B cell KO reduces PCL-mediated fibrosis. (**A**) Histological staining of 6-week PCL implants in BL6-WT and BKO with Masson’s Trichrome for collagen fibrils. Scale bar = 400µm. (**B**) Gene expression analysis of fibrosis related genes including col1a1, S100a4, p21, TGF-B, IL-23a, Col3a1 via qRT-PCR. (**C**) Immunofluorescence staining of CD20^+^ cells (yellow) and HLA-DR^+^ (red) in human tissue samples surrounding breast capsule implants. Scale bar = 100µm. Data are mean ± SD, n=3 for NT controls, 4 for PCL biomaterial, Two-way ANOVA with subsequent multiple comparison testing [C: **p<0.001, **p<0.01, *p<0.05].

Finally, to determine potential clinical relevance we assessed the potential role of B cells in FBR to human breast implant tissue expanders which have been previously demonstrated to exhibit a FBR and Type 3 (IL-17-mediated response) (*4, 23*). Small clusters of CD20^+^ B cells (Fig. 4C) were present in tissue adjacent to the implant. These B cells were also HLA-DR^+^, suggesting the chronic presence of antigen presenting B cells in the capsule around clinical breast implants (Fig. 4C).

## Discussion

The innate immune response to tissue damage and biomaterials is well recognized, along with a tissue specificity to this behavior. Tissue damage signals after injury activate multiple cell types through multiple pathways, including neutrophils, eosinophils, monocytes, macrophages, among other immune cells. Neutrophils can orchestrate and prime adaptive immune cell responses through secretion of chemokines/cytokines and the release of Neutrophil Extracellular Traps (*36–38*). Chronic neutrophilia is also associated with Type 3 (IL-17 immune responses) (*39*). Eosinophils are of particular interest in the realm of natural biomaterials as these cells can prime the local tissue response and stimulate an IL-4/Type 2 immune response that can contribute to wound healing (*40, 41*). Monocytes and macrophages are also known to alter their immune phenotypes along a wide spectrum of pro-inflammatory and pro-resolution outcomes in response to biomaterials (*42–45*). Their phenotypes are influenced by numerous factors including the tissue-specific immune environment and biomaterial physical-chemical properties that direct tissue healing and repair (*46–48*). The paradigm of the FBR to biomaterials has traditionally focused on these innate immune cells and their impact on fibrosis. However, there is growing recognition of the adaptive immune response to tissue damage and repair. For example, Ramos and colleagues found that antigen-specific T cells recognizing tissue-specific extracellular matrix molecules developed after cardiac injury (*49*). In the case of CNS injury, Kipnis and colleagues discovered that MHCII was required for Th2-mediated repair (*50*). Tissue damage can occur with biomaterial implantation and tissue remodeling suggesting that an adaptive immune response may also occur (*22, 23*). T cell responses to both synthetic and naturally-derived biomaterials has now been demonstrated in multiple studies (*22, 23*).

B cells are known to participate in the immune response to tissue damage and wound healing through antibody production and cytokine secretion (*51–53*). Following injury, IgM and IgG antibodies rapidly increase in the tissue and blood, where they accelerate critical functions in healing by binding to wounded tissues, inducing phagocytosis by Fc receptors on neutrophils or macrophages (*51, 53*). Importantly, autoantibodies are present in the injury site and connect both the local injury immune response and splenic responses, as splenectomies result in prolonged presence of neutrophils and macrophages in an injury site (*53*). Multiple studies have defined direct roles for B cells in repair of skin excisions and bone fractures along with repair of kidney damage and intestinal injury (*16, 54–57*). The addition of B220^+^CD19^+^ B cells directly on to diabetic wound skin excisions significantly accelerated neutrophil invasion, decreased apoptosis in the wound bed and reduced time to skin closure (*16*). Regulatory B10 cells, which secrete IL-10, mediate intestinal inflammation and orchestrate bone fracture repair in a time-sensitive manner (*56*).

B cell research in the context of medical implants historically focused on antibody production. In particular, the concerns over deleterious responses to breast implants in the 1990s prompted multiple safety studies (*58–60*). These studies of breast implant safety primarily investigated antibody production, specifically antibodies that might recognize the implant itself. They found that patients who had received silicone elastomer implants had elevated anti-polymer reactive antibodies in the serum, which were non-existent in non-implanted patients. Further studies have disputed this finding, suggesting that antibody reactivity was to the hydrophobic nature of the silicone and not-implant specific (*61*). Review of the safety studies by the US Institute of Medicine in 2000 led to an inconclusive finding, and silicone implant use has since continued in the clinic (*62*). While ongoing work seeks to assess the presence and antigen-specificity of antibodies, B cells in wound healing are not limited to antibody production. The functions of B cells, the generation of germinal centers in response to wounds, and the response of the splenic B cells can also highlight additional B cell contributions to wound healing and to the biomaterial response.

Biomaterial scaffolds have previously been shown to induce B cells to distinct phenotypes and functions. Polymeric nanoparticles and vaccines serve both as delivery vehicle and an adjuvant to enhance recognition and antibody production. Babensee and colleagues found that polymeric scaffolds serve a deleterious adjuvant function that enhances recognition of transplanted cells that decreases cell viability and regenerative function (*63*). In this work, we investigated the B cell response to a biological ECM scaffold and a synthetic polyester. ECM-based materials contain proteins that serve as obvious antigens to an adaptive material response (*6*). Furthermore, Allman and colleagues presented systemic TH2 activity with ECM exposure, confirming that both arms of the adaptive immune system are activated with these materials (*6*). These studies support our results that show plasmablast development in B cells exposed to SIS-ECM material. Mice treated with ECM after injury demonstrated elevated levels of IgG1 and IgM at 6 weeks, which has previously been demonstrated to facilitate wound healing in a clinical model of skin regeneration (*53*). This introduces a novel perspective of B cell response to biomaterials, promoting the classification of both the antibodies and B cell subsets found in wounds. Lack of B cells did not appear to impact the response to ECM, although tissue repair was not studied in detail.

In contrast to ECM-based biomaterials, synthetic polymers such as PCL do not have an obvious antigen for MHC presentation. While we previously demonstrated an antigen specific response to PCL (*23*), the source of this antigen remains unknown. Results from this study highlight PCL implantation increases antigen presentation by B cells. B cells have a unique capacity for antigen presentation that has been implicated in infection, disease pathogenesis and autoimmunity (*64–66*). Specifically, B cells present specific antigens to activate other B cells and cognate T cells. The specific presentation of antigens can induce T cell proliferation and has been seen to promote a Type 2 wound healing response in chronic allergic lung diseases. B cells are also capable of non-specific antigen presentation, which results in inactivation of T cells or induction of regulatory T cell differentiation (*67*). The efficacy and efficiency through which B cells serve as antigen presenting cells is tightly regulated, as MHC-II presentation is dependent on B cell developmental stage. As with all things in immunology, the exact role of B cells as antigen presenting cells is context dependent. However, it is known that B cell antigen presentation is a highly coordinated uptake which induces cell polarization and usually occurs in the secondary lymph organ.

Splenic B cells also expressed genes associated with inflammation in response to PCL implantation, such as *S100a8*, suggesting that B cells are participating in the FBR beyond antigen presentation. Lack of B cells in the MuMt^−^ mice appeared to reduce fibrosis around the materials further supporting the role of B cells in the FBR fibrotic response. While we only studied two materials in this paper, they represent two very distinct potential B cell responses to biomaterials. They were selected both as examples of different compositions but also producing distinct immune responses. Other materials will likely produce responses that fall somewhere in the middle on this wide spectrum of material properties and responses.

Traditionally, the foreign body response to biomaterial implants and scaffolds has been considered primarily a local phenomenon. The involvement of adaptive immune cells, such as T cells and B cells, in the FBR and tissue repair process opens the door to considering a systemic immune response. Here, we found changes in B cell expression profiles in the local draining lymph nodes and the spleen. Gene expression patterns in the splenic B cells shifted with injury alone and further differentiated depending on the biomaterial composition. While B cells may be small at the local injury or implant site, they have physiological impact, particularly in the case of synthetic materials. The potential interplay between a local implant and systemic factors introduces the concepts that a biomaterial may impact the overall physiological state of an organism, and vice versa, the physiological state of the organism may impact the response to injury, implants and scaffold repair strategies.

## Materials and Methods

### Surgical injury and material implantation

All animal procedures performed with approval by Johns Hopkins University Institutional Animal Care and Use Committee. The animals were all female mice aged 6 to 9 weeks old. Two strains were used: wild-type C57BL/6J (The Jackson Laboratory, stock #00064) and muMt^−^ (B cell KO) on C57BL/6J background (The Jackson Laboratory, stock #002288). Bilateral traumatic muscle defects were created as previously described (*22, 23, 68*). While under anesthesia, a 3 mm x 3mm x 4 mm section of quadriceps femoris tissue was excised from both hindlimbs. Following injury, the quadriceps muscle defects were filled directly with 30 mg of a synthetic (PCL, particulate, Mn = 50,000 g/mol; mean particle size <600μm, Polysciences) or biological (ECM, decellularized porcine small intestine submucosa) scaffold material. For control surgeries (considered no-implant with injury controls) 0.05 ml of phosphate-buffered saline (PBS) was dispensed directly into the wound. All materials were disinfected with UV before use. Immediately after surgery, mice were given subcutaneous carprofen (Rimadyl, Zoetis) at 5 mg/kg for pain relief. Mice were euthanized for sample harvests at Day 3, Day 5, 1,3, and 6 weeks after surgery.

### Sample collection and digestion for single cell analyses

Murine tissue samples (quadriceps femoris muscle and inguinal lymph nodes) were obtained for single cell flow cytometry and/or fluorescence activated cell-sorting. Quad tissue samples were finely diced and digested with 1.67 Wünsch U/mL Liberase TL (Roche Diagnostics) and deoxyribonuclease I (0.2 mg/ml; Roche Diagnostics) in RPMI-1640 medium (Gibco) for 45 min at 37°C. The digested quad tissues were ground through 70-µm cell strainers (Thermo Fisher Scientific), rinsed with RPMI-1640 + 5% fetal bovine serum. Percoll (GE Healthcare) density gradient centrifugation was used to remove debris from quad digestions and enrich the leukocyte fraction. Inguinal lymph node samples were ground through 70-µm cell strainers with excess RPMI-1640 + 10 mM HEPES buffer solution. The enriched single-cell suspensions were washed and stained following antibody panels **Tables S1-S3**, respective to the intended application. No differences in single-cell isolation from different material environments were observed.

### qRT-PCR assay

Isolated tissues were digested using TRIzol reagent to isolate total RNA. RNeasy Mini-Kit (Qiagen) purified mRNA from total RNA. qRT-PCR was performed using TaqMan Gene Expression Master Mix (Applied Biosystems) consistent with the manufacturer’s instructions. In brief, 2µg of total RNA was used to synthesize complementary DNA (cDNA) using SuperScript IV VILO Master Mix (Thermo Fisher Scientific). The cDNA concentration was set, according to manufacturer recommendation, at 100 ng/well (in a total PCR volume of 20-µl). qRT-PCR assays were performed as TaqMan single-plex assays on the StepOne Plus Real-Time PCR System (Thermo Fisher Scientific), using the manufacturer recommended settings for quantitative and relative expression. For tissue samples, Hprt, Rer1 or Oaz1 were used as the reference gene and samples were normalized to either no-treatment or injury with saline depending on application. All qRT-PCR data were analyzed using the Livak method, where ΔΔCt values are reported as relative quantification (RQ) values calculated by 2-ΔΔ Ct. RQ values are represented by geometric means, with geometric standard deviation represented by error bars. Low expressing mRNA transcripts were preamplified using 250 ng of cDNA and the TaqMan Pre-Amp System (Thermo Fisher Scientific) with 14 cycles of amplification according to manufacturer recommendation. Primer probes used for qRT-PCR and preamplification listed in **Table S4**.

### Flow cytometry and fluorescence activated cell-sorting

For cell isolation using fluorescence-activated cell sorting (FACS), the single cell suspensions from digested quad and inguinal lymph node samples were stained for 30 min at 4°C using Viability Dye eFluor 780 (eBioscience). For intracellular staining, the cells were stimulated with Cell Stimulation Cocktail (plus protein transport inhibitors) (eBioscience) diluted in complete culture media IMDM (Iscove’s modified Dulbecco’s medium) supplemented with 5% fetal bovine serum for 4 hours. Cells were then washed, surface-stained, fixed/permeabilized with Cytofix/Cytoperm (BD), and then stained for intracellular markers. Flow cytometry was conducted using Attune NxT Flow Cytometer (Thermo Fisher Scientific). Antibody panels are listed in **Tables S1-S3**.

### NanoString gene expression analysis

B cells were isolated from inguinal lymph node samples as previously described and stained for CD19 and CD3 (**Table S3**). CD19^+^ cells were sorted using FACS and mRNA was then isolated. After lysis in RLT plus buffer containing 2-mercaptoethanol, the RNA was purified using the RNeasy Plus Micro Kit (Qiagen) and then quantified using a Qubit RNA fluorometric assay high-sensitivity kit (Thermo Fisher Scientific). The gene expression analysis for n =3 biologically independent samples was conducted with the nCounter® PanCancer Mouse Immune Profiling Panel (XT-CSO-MIP1-12, NanoString Technologies). Based on mRNA quantification, 50 ng of mRNA per sample was added to a barcoded probe set reagent and hybridized for 20 hours at 65°C according to the manufacturer’s protocol. NanoString data were processed with the nSolver 4.0 software kit according to the manufacturer’s protocol. Differentially expressed genes (P <0.05 and positive fold change) for each sample group were used for further analysis.

### Single cell RNA sequencing

The 10x Chromium instrument with 5’ v2 chemistry was used to generate single cell RNA sequencing libraries. Directly prior to loading in the chip, hashed samples were pooled in equal amounts. Encapsulation and library prep were performed according to manufacturer’s recommendations. Briefly, cells and beads are encapsulated in water droplets in a water-in-oil emulsion. Reverse transcription was then completed, tagging cell RNA and hashing antibodies with bead-specific oligonucleuotide barcodes. We then ran whole transcriptome amplification and separated the hashing antibody cDNA from the transcript cDNA. A sequencing library was generated for each portion. Finally, we preferentially amplified B cell receptor from an aliquot of the transcript cDNA and prepared a separate library for VDJ sequencing. Transcript and hashing libraries were pooled at a ratio of 9:1. The pooled libraries were sequenced to a depth of 50,000 reads per cell on the transcript library. The VDJ libraries were sequenced at 100,000 molecule reads per cell. In both cases 10x recommended sequencing configurations were followed.

### Transcriptome read alignment and processing

Read alignment for all three libraries was performed using Cellranger 3.0 recommended settings and the provided mm10 references where applicable. Seurat was used for processing steps where other software packages are not specified. Genes expressed in fewer than 0.1% of cells were removed and cells with less than 500 UMIs were removed. HTODemux was used with the hashing data to group cells into condition as well as remove condition-doublets. Subsequent counts were log-normalized. Cell cycle scores were calculated using Seurat’s CellCycleScoring with previously defined gene sets (*69*) and the total percent of mitochondrial genes calculated. Normalized gene expression values were scaled with linear regression of genes on Cell cycle score, total UMI count, and percent mitochondrial gene contribution.

### PCA, dimensional reduction, clustering, and differential expression

Principle component analysis was performed on the scaled data and the top 50 used for downstream analysis. UMAP was used for dimensional reduction and visualization. A shared nearest neighbor graph was constructed from the principle components and subsequent Louvain clustering used to identify clusters. Mann-Whitney U tests were used to test differential expression of each cluster in comparison to all other cells in the data set. The top 10 genes by fold-change by cluster were used to generate a heatmap of differentially expressed genes.

### VDJ library processing

VDJ librarys were processed with Cellranger 3.0’s VDJ function with the provided mm10 references. Cellranger’s clonotype barcodes were then cross-referenced with hashing barcodes to label each cell for condition and singlet status. The resulting clonotype tables are reported (Fig. S4).

### Serum/Plasma (ELISAs)

Blood serum samples were collected by submandibular bleeding and diluted 1:20,000 using Abcam Mouse IgG1, IgM, IgG2a ELISA Kits (catalog no. ab133045, ab133047, ab133046, respectively). ELISA was performed according to the manufacturer recommended protocol. Briefly, the PCL-coated plate was blocked with the assay diluent for 1 hour before loading. Each serum dilution was loaded into the plate after blocking and incubated for 2 hours. After washing, anti-mouse IgG1, IgM, or IgG2a was added to conjugate the bound antibody for 1 hour. After washing, 30 minute incubation with provided Substrate solution, and addition of provided Stop Solution, absorbance was read at 450 nm with correction between 570 and 590 nm.

### Mouse samples for histopathology

Upon tissue harvest, immediate fixation was performed in 10% neutral buffered formation (48hr), followed by step-wise dehydration to ethanol (200 proof). Tissues were then cleared with xylenes, and paraffin embedded. Sectioning of samples was performed with a Leica RM2255 microtome in a transverse orientation (7 µm thickness). Histopathological examination was conducted with Masson’s trichrome (Sigma-Aldrich) according to manufacturer protocols. Brightfield imaging of stained slides was performed.

### Human samples for histology

Tissue was acquired from patients undergoing breast implant exchange or replacement surgeries (n = 12, Johns Hopkins University Institutional Review Board exemption IRB00088842); the breast implants were surgical discards that were deidentified. Each section was weighed, and 0.25 g of tissue was dissected for histology. Peri-implant samples included tissues surrounding implants with both smooth and textured surface properties. All implants had a silicone shell and were either temporary tissue expanders filled with saline or air or permanent implants filled with silicone or saline. Average patient age was 56 years old (range of 41 to 70 years old), and the average implant residence time was 41 months (range of 1 to 360 months).

Tissues were harvested and fixed in 10% neutral buffered formalin for 24 hours before dehydration in EtOH, cleared with xylene, and embedded in paraffin. Samples were sectioned as 7 μm slices using the Leica RM2255 microtome. Samples were baked, deparaffinized, and re-fixed in neutral buffered formalin. Antigen retrieval was achieved by heating slides in 10mM sodium citrate. Endogenous peroxidases were quenched using 3% H2O2 (Sigma-Aldrich) and aldehydes were quenched in glycine. Samples were incubated with 10% bovine serum albumin (BSA) (Sigma-Aldrich), and 0.05% Tween solution. Samples were incubated with anti-cD20 antibody (Abcam ab78237) at 1:500 dilution for 10 min in 10%BSA. Samples were washed and underwent Opal HRP polymer incubation (Akoya Biosciences, ARH1000EA) for 10 min. Samples were washed again and underwent incubation with Opal 570 fluorophore in 1x amplification diluent (Akoya Biosciences) for 10 min. Samples underwent final washing and staining with DAPI stain for 5min. Coverslips were applied and slides were imaged using a Zeiss Axio Observer with Apotome.2 and Zeiss Zen Blue software version 2.5.

## Supporting information

Data files S1. Multiplex gene expression analysis of B cells sorted from the ILN 1 week after injury.

## General

We acknowledge S. Ganguli for assistance with NanoString multiplex gene expression platform.

## Author contributions

E.M.M., and J.H.E. conceptualized the study. E.M.M. and J.H.E. discussed and formulated experimental design for the study. E.M.M., D.R.M., and C.C. contributed to analysis and interpretation of results. E.M.M., D.R.M., J.A.G., and H.Y.C. contributed to conducting experimental procedures and assisted with the harvest/preparation of tissue for flow cytometry. A.T and S.G. assisted with flow cytometry for cell sorting and Nanostring gene expression analysis. J.A.G. assisted with histology staining and imaging. G.D.R. provided human surgical discard samples. L.D.H. discussed and assisted in interpreting the results and commented on the manuscript. E.M.M. and J.H.E. wrote the manuscript with input from all co-authors.

## Data and materials availability

All data needed to evaluate the conclusions in the paper are present in the paper and/or the Supplementary Materials. Additional data related to this paper may be requested from the authors.

## Supplementary Materials

(Data File IN ATTACHMENT)

**Data files S1. Multiplex gene expression analysis of B cells sorted from the ILN 1 week after injury**.

**Fig S1.**
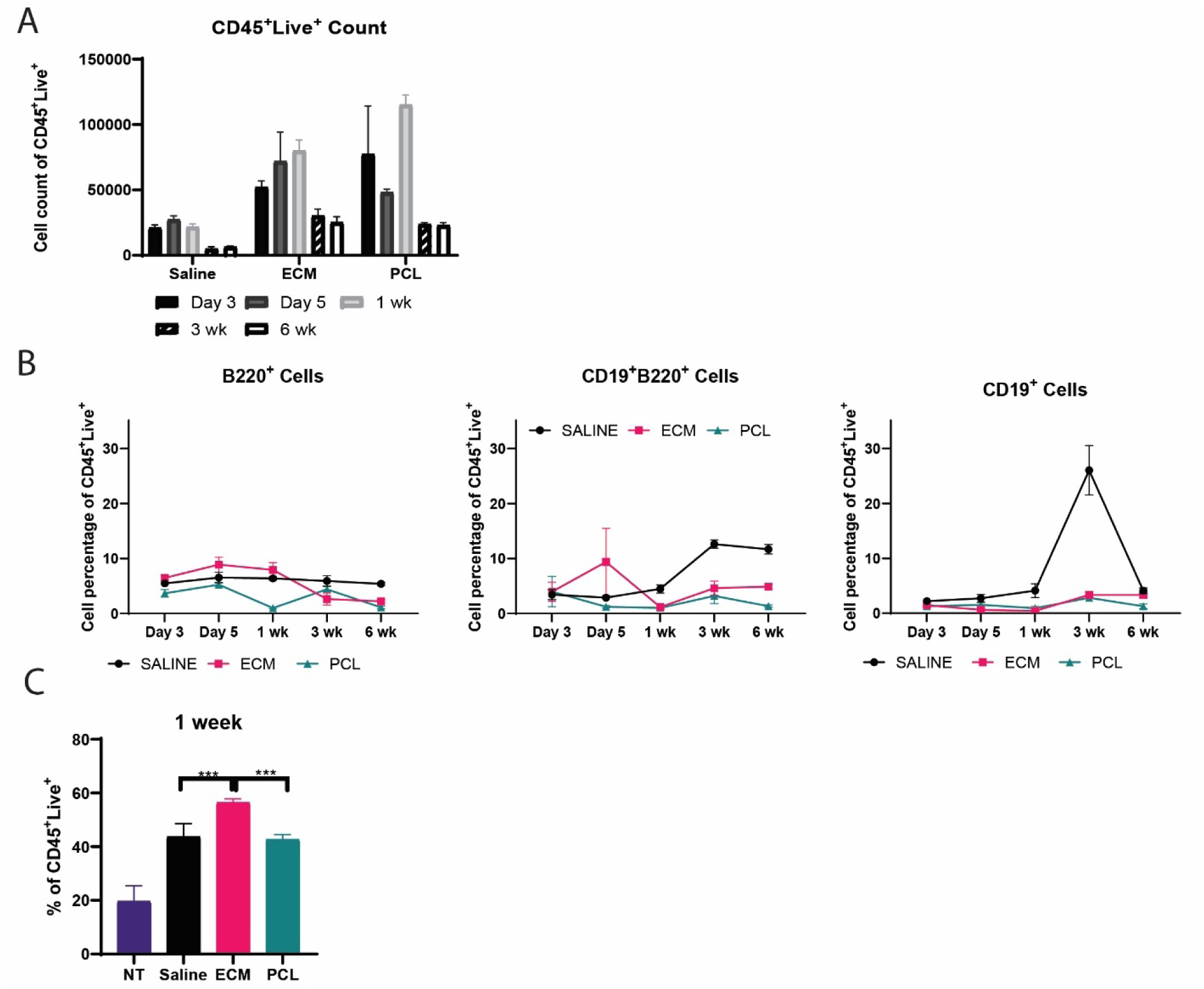
Total immune and B cell content in quad tissue and draining inguinal lymph node after injury with or without biomaterial implant. (**A**) CD45^+^ immune cells present in injured quad tissue over time. (**B**) Evolution of CD19^+^B220^−^, CD19^+^B220^+^, CD19^−^B220^+^ B cells as a percentage of the CD45^+^ cells in the quad over time. (**C**) B220^+^CD19^+^ B cells in ILN 1 week post injury. Data are mean ± SD, n=4, Two-way ANOVA with subsequent multiple comparison testing [C: **p<0.001, **p<0.01, *p<0.05].

**Fig S2.**
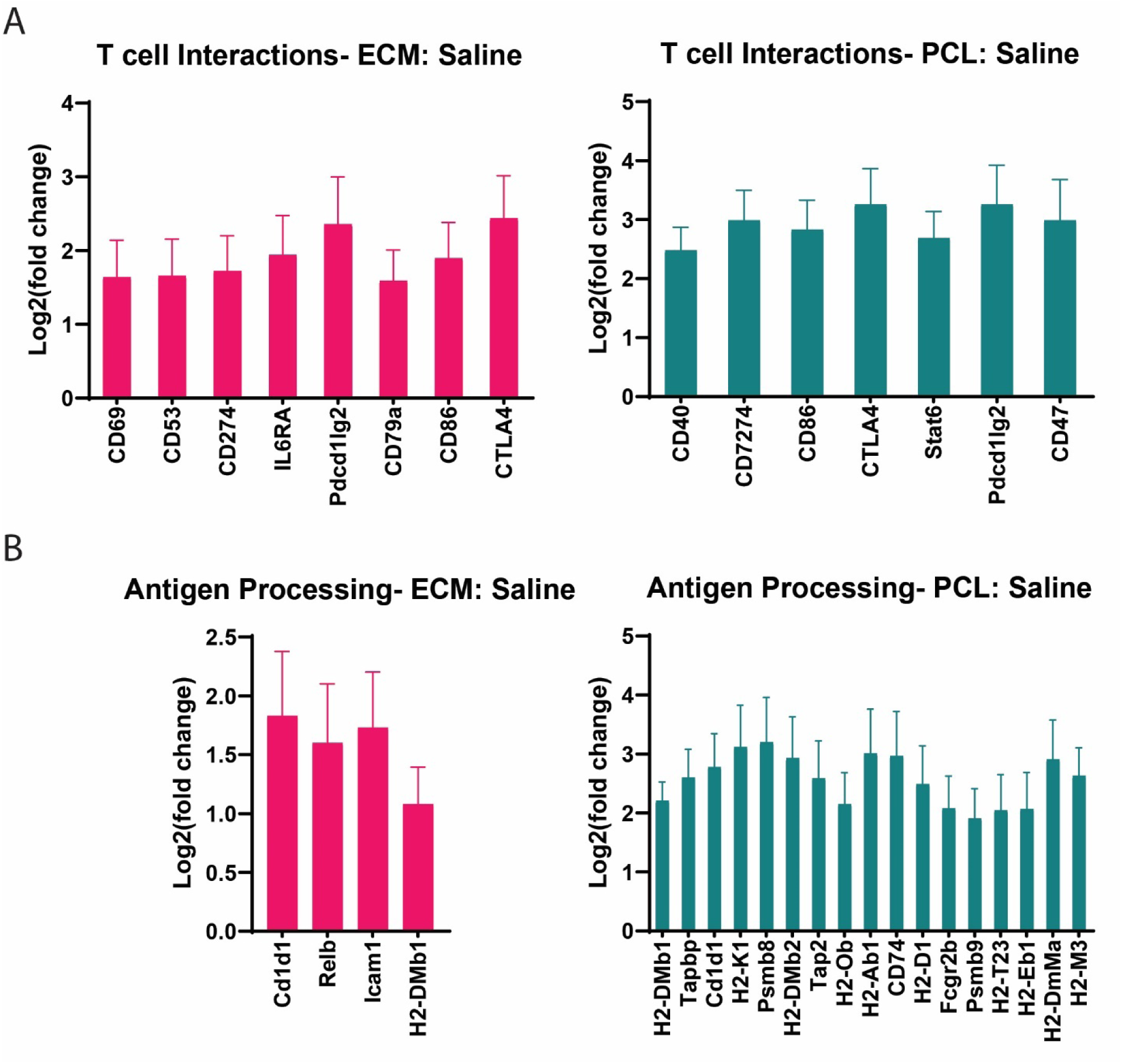
Gene expression related to T cell interactions and antigen processing in B cells from the ILN at 1 week. (**A-B**) ECM and PCL treatment groups present upregulated genes associated with (**A**) B and T cell interactions and (**B**) antigen presentation when compared to Saline treated animals.

**Fig S3.**
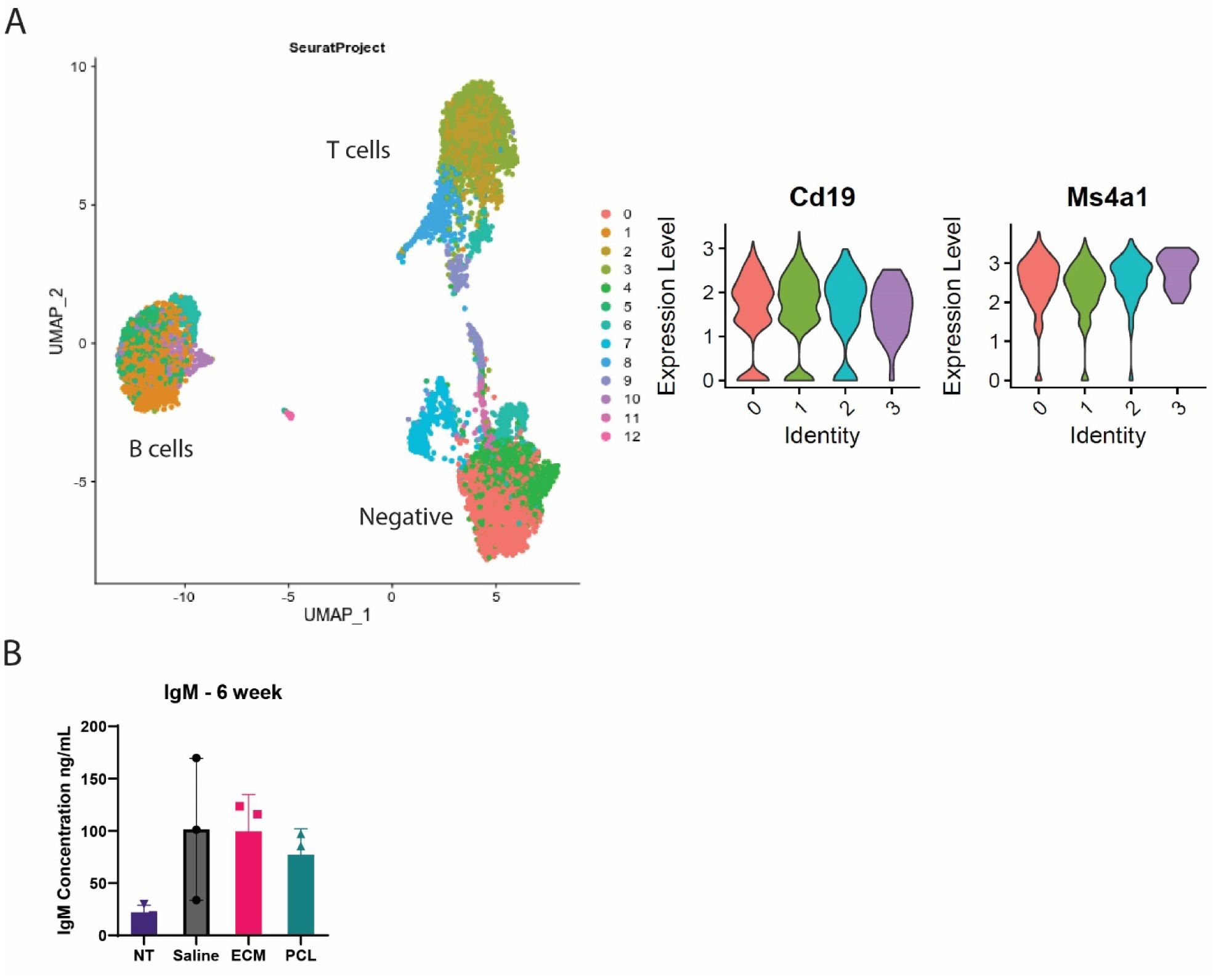
Identification of B cell local and systemic response. **(A)** A uniform manifold approximation and projection (UMAP) of sorted B and T cells from ILN at 3 weeks. Differentiation of cells based on CD19 and Ms4a1. **(B)** IgM serum levels increased in each condition at 6 weeks post injury.

**Fig S4.**
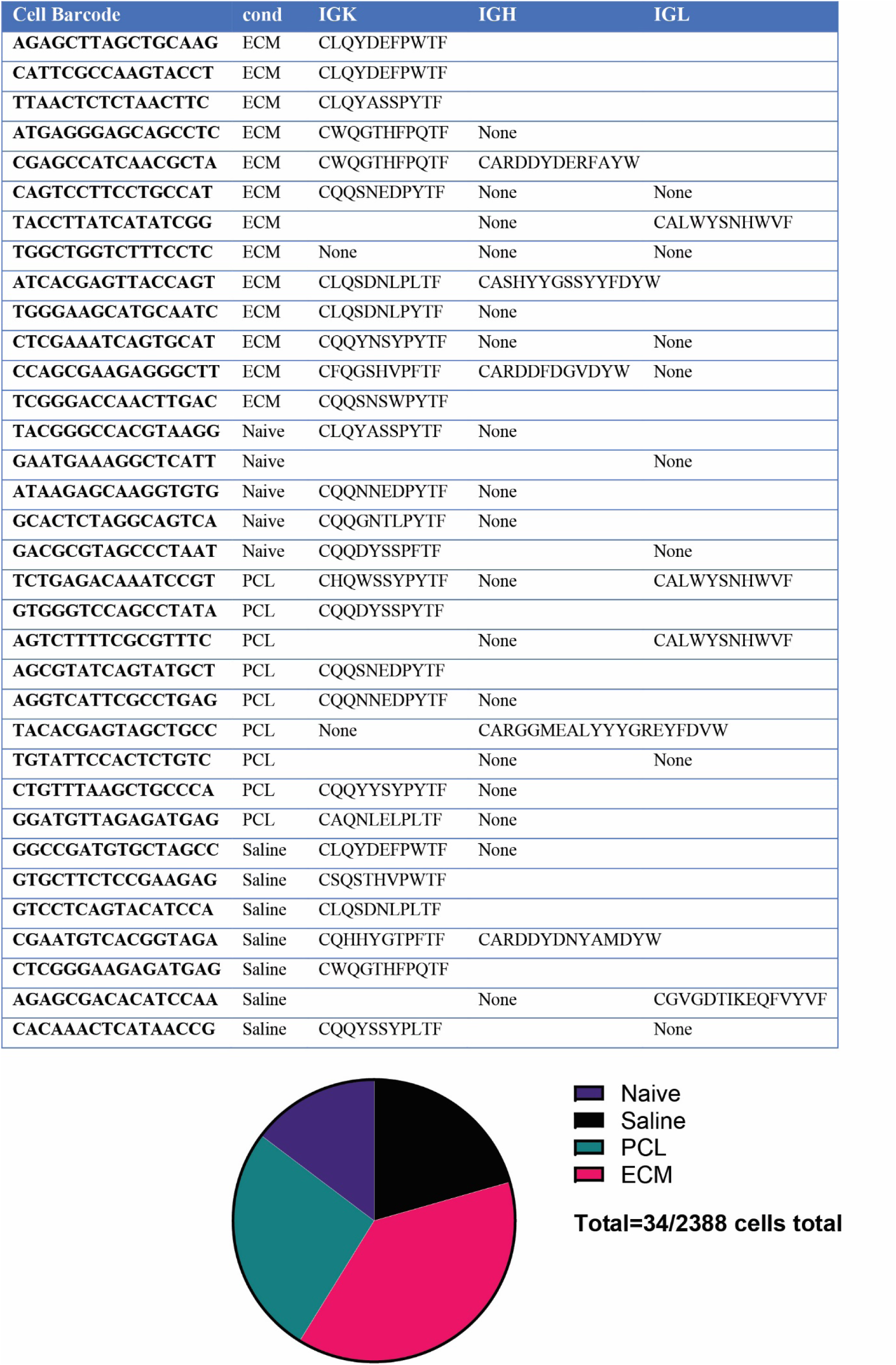
VH Repertoires identified from single cell RNA sequencing. Clonal cells are shown by condition and clonal sequence. Pie plot shows number of clonal cells by condition.

**Fig S5.**
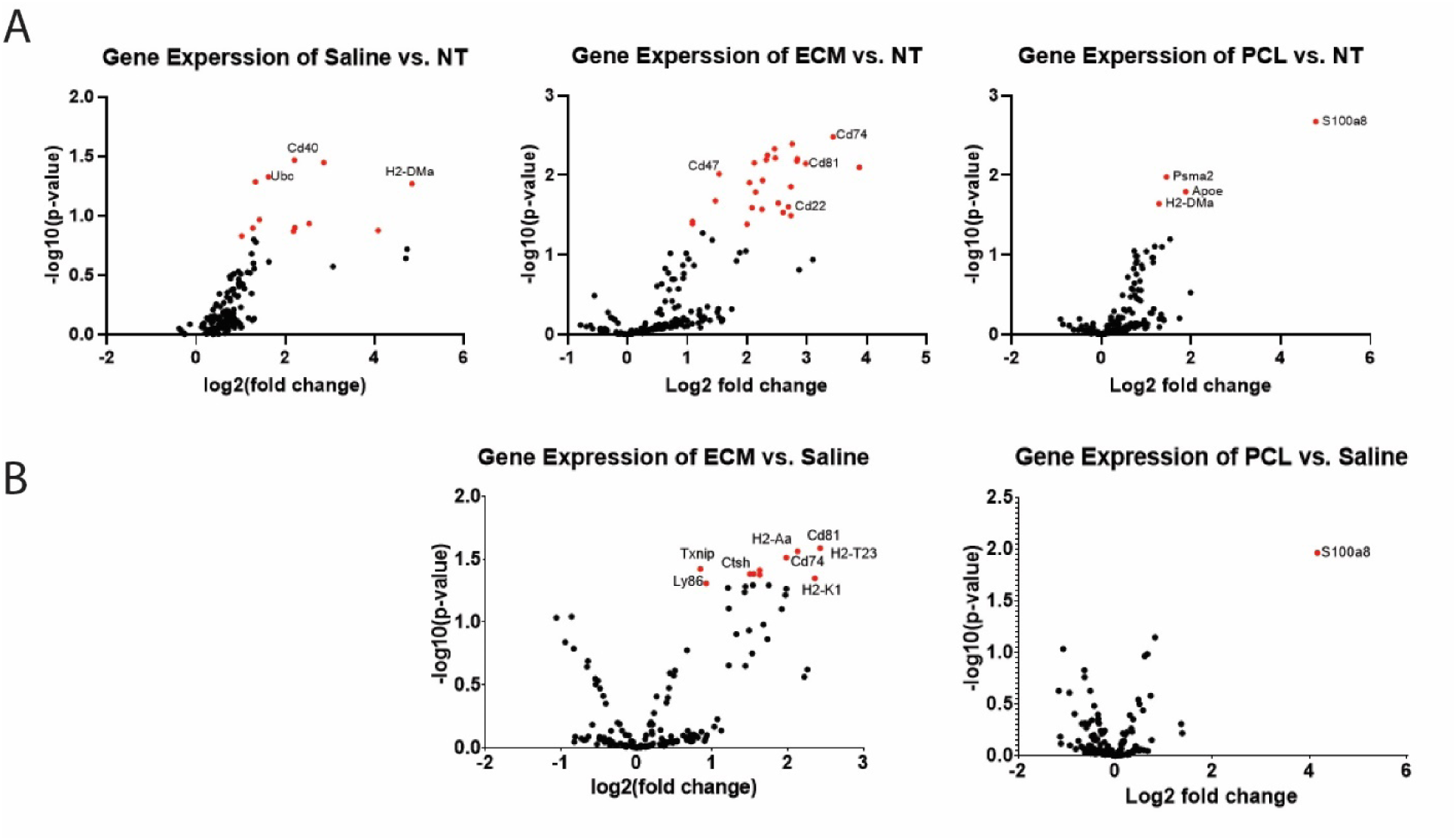
Splenic B cells gene expression. **(A)** Gene expression volcano plots of expressed genes compared to no treatment (NT). **(B)** Gene expression volcano plots of ECM and PCL compared to Saline. Red indicates statistically significance compared to NT, *p<0.05 in A, compared to Saline in B.

**Fig S6.**
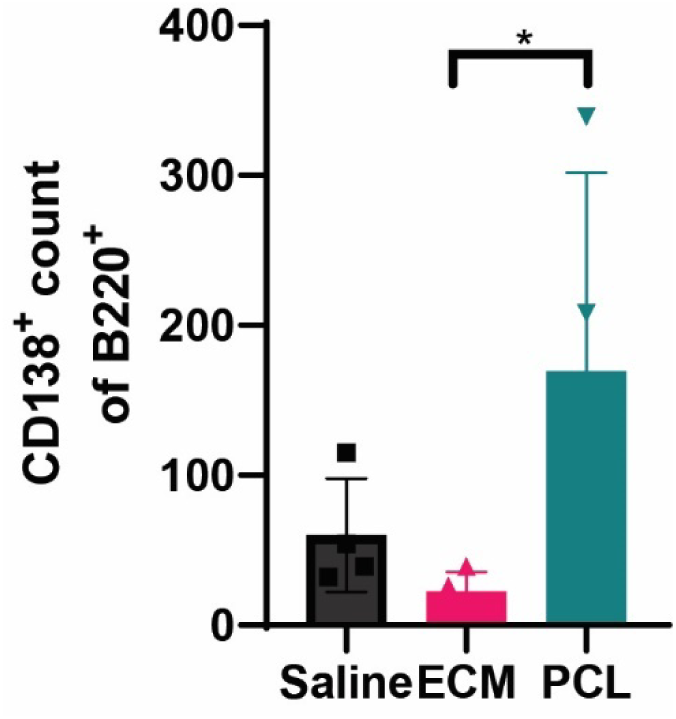
PCL induces CD138^+^ B220^+^ cells in the quad tissue. CD138^+^B220^+^ cell population in quad tissue at 3 weeks post-injury. Data are means ± SD, n=4, Two-way ANOVA with subsequent multiple comparison testing [*p<0.05].

**Fig S7.**
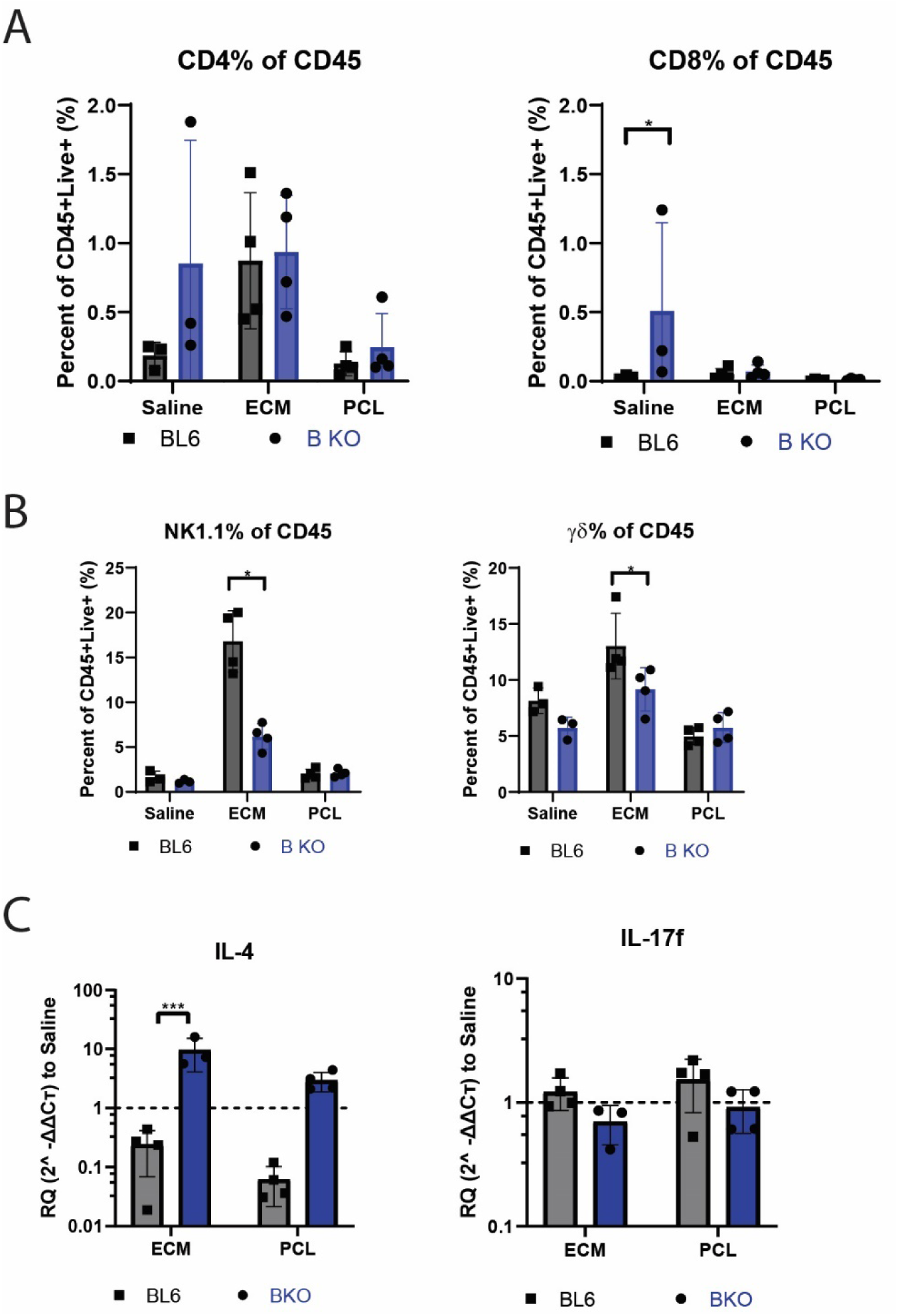
Cell and genetic differences between MuMt^−^ mice and BL6 mice. **(A-B)** Flow cytometry analysis of (A) CD4 and CD8 T cells and (B) NK1.1 and γγγγTcells as a percentage of CD45^+^Live^+^ cells in quad at 1-week post-injury. **(C)** Gene expression analysis of *il4* and *il17f* at 6-weeks post injury. Data are means ± SD, n=4, Two-way ANOVA with subsequent multiple comparison testing [*p<0.05].

**Table S1.** B cell flow cytometry staining panel.

| Table S1. Antibodies for B cell flow cytometry panel |  |  |  |  |
| --- | --- | --- | --- | --- |
| Fluorophore | Marker | Clone | Catalog | Manufacturer |
| Brilliant Violet 421™ | CD19 | 6D5 | 115537 | Biolegend |
| BV605 | CD45 | 30-F11 | 103155 | Biolegend |
| BV711 | CD11b | M1/70 | 101242 | Biolegend |
| AF488 (FITC) | CD3 | 17A2 | 100210 | Biolegend |
| PE | CD138 | 281-2 | 142503 | Biolegend |
| PE/Dazzle 594 | I-A/I-E | M5/114.15.2 | 107648 | Biolegend |
| PE-Cy7 | B220 | RA3-6B2 | 103221 | Biolegend |
| APC | CD27 | LG-3A10 | 124211 | Biolegend |
| AF700 | CD5 | 53-7.3 | 100635 | Biolegend |
| eFluor780 | Viability |  | 65086514 | ThermoFisher |

**Table S2.**
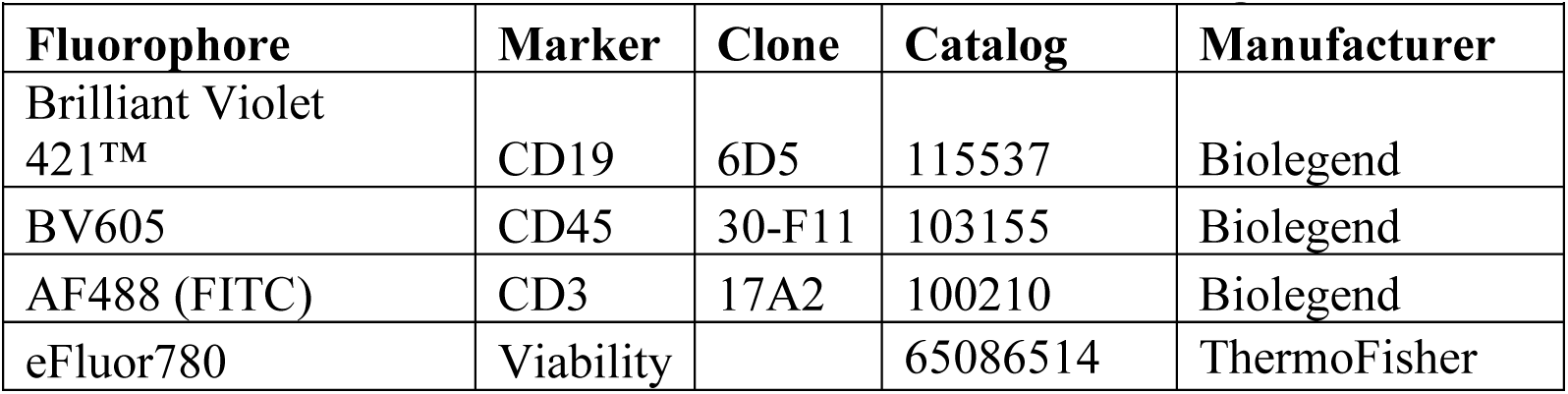
B cell sorting antibodies for Fluorescent Assorted Cell Sorting.

**Table S3.** Flow cytometry antibody panel for T cell characterization.

| Table S3. Antibodies for flow cytometry (T cell panel) |  |  |  |  |
| --- | --- | --- | --- | --- |
| Fluorophore | Marker | Clone | Catalog | Manufacturer |
| BV421 | FoxP3 | MF-14 | 126419 | Biolegend |
| V500 | CD45 | 30-F11 | 561487 | BD Biosciences |
| BV605 | NK1.1 | PK136 | 108739 | Biolegend |
| BV711 | CD8 | 53-6.7 | 100748 | Biolegend |
| AF488 (FITC) | CD3 | 17A2 | 100210 | Biolegend |
| PerCP-Cy5.5 | CD19 | 6D5 | 115534 | BioLegend |
| PE | IL-4 | 11B11 | 504103 | Biolegend |
| PE-Dazzle 594 | I-A/I-E | M5/114.15.2 | 107648 | Biolegend |
| PE-Cy7 | CD4 | GK1.5 | 100422 | Biolegend |
| APC | IFNγ | XMG1.2 | 505810 | Biolegend |
| AF700 | IL-17a | TC11-18H10.1 | 506914 | Biolegend |
| eFluor780 | Viability |  | 65086514 | ThermoFisher |

**Table S4.**
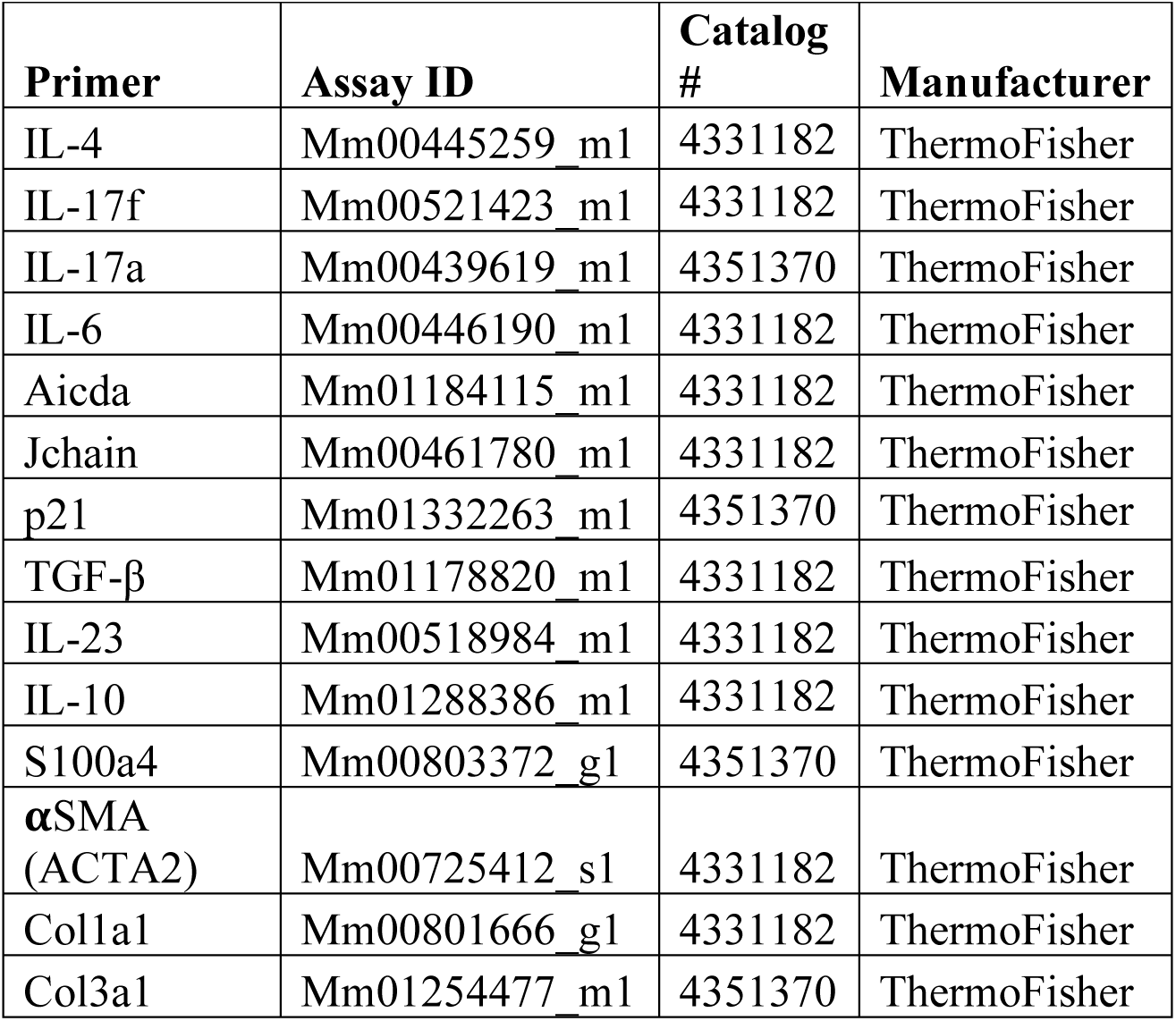
Murine TaqMan gene expression primers.

| **Table S4. Murine TaqMan gene expression primers** |  |  |  |
| --- | --- | --- | --- |
| **Primer** | **Assay ID** | **Catalog #** | **Manufacturer** |
| IL-4 | Mm00445259_m1 | 4331182 | ThermoFisher |
| IL-17f | Mm00521423_m1 | 4331182 | ThermoFisher |
| IL-17a | Mm00439619_m1 | 4351370 | ThermoFisher |
| IL-6 | Mm00446190_m1 | 4331182 | ThermoFisher |
| Aicda | Mm01184115_m1 | 4331182 | ThermoFisher |
| Jchain | Mm00461780_m1 | 4331182 | ThermoFisher |
| p21 | Mm01332263_m1 | 4351370 | ThermoFisher |
| TGF-β | Mm01178820_m1 | 4331182 | ThermoFisher |
| IL-23 | Mm00518984_m1 | 4331182 | ThermoFisher |
| IL-10 | Mm01288386_m1 | 4331182 | ThermoFisher |
| S100a4 | Mm00803372_g1 | 4351370 | ThermoFisher |
| αSMA<br>(ACTA2) | Mm00725412_s1 | 4331182 | ThermoFisher |
| Col1a1 | Mm00801666_g1 | 4331182 | ThermoFisher |
| Col3a1 | Mm01254477_m1 | 4351370 | ThermoFisher |

## References

1. J. Couzin-Frankel, Cancer immunotherapy. Science (80-.). 342, 1432–1433 (2013).

2. L. Steinman, Immune therapy for autoimmune diseases. Science (80-.). 305, 212–216 (2004).

3. L. Chung, D. R. Maestas, F. Housseau, J. H. Elisseeff, Key players in the immune response to biomaterial scaffolds for regenerative medicine. Adv. Drug Deliv. Rev. 114, 184–192 (2017).

4. D. Wolfram, E. Rabensteiner, C. Grundtman, G. Böck, C. Mayerl, W. Parson, G. Almanzar, C. Hasenöhrl, H. Piza-Katzer, G. Wick, T Regulatory Cells and TH17 Cells in Peri–Silicone Implant Capsular Fibrosis. Plast. Reconstr. Surg. 129, 327e – 337 (2012).

5. S. B.M., P. R. J., D. C.L., W. M.T., A. F., B. M., T. N.J., W. D.J., S. T.W., W. A., B. E.H.P., D. J.L., F. L.E., B. S., B. S.F., An acellular biologic scaffold promotes skeletal muscle formation in mice and humans with volumetric muscle loss. Sci. Transl. Med. 6 (2014) (available at http://stm.sciencemag.org/content/6/234/234ra58.full.pdf&5Cn http://ovidsp.ovid.com/ovidweb.cgi?T=JS&PAGE=reference&D=emed12&NEWS=N&AN=2014313752).

6. A. J. Allman, T. B. Mcpherson, S. F. Badylak, L. C. Merrill, B. Kallakury, C. Sheehan, R. H. Raeder, D. W. Metzger, Xenogeneic extracellular matrix grafts elicit a TH2-restricted immune response. Transplanation. 71, 1631–1640 (2001).

7. T. W. Gilbert, T. L. Sellaro, S. F. Badylak, Decellularization of tissues and organs. Biomaterials. 27, 3675–3683 (2006).

8. A. Singh, Biomaterials innovation for next generation ex vivo immune tissue engineering. Biomaterials. 130, 104–110 (2017).

9. M. Rahmati, C. P. Pennisi, E. Budd, A. Mobasheri, M. Mozafari, in Cell Biology and Translational Medicine, Volume 4: Stem Cells and Cell Based Strategies in Regeneration, K. Turksen, Ed. (Springer International Publishing, Cham, 2018; https://doi.org/10.1007/5584_2018_278), pp. 1–19.

10. J. M. Anderson, A. Rodriguez, D. T. Chang, Foreign body reaction to biomaterials. Semin. Immunol. 20, 86–100 (2008).

11. J. M. Anderson, A. K. McNally, Biocompatibility of implants: lymphocyte/macrophage interactions. Semin. Immunopathol. 33, 221–233 (2011).

12. J. M. Anderson, Inflammatory response to implants. ASAIO J. 34, 101–107 (1988).

13. K. Murphy, C. Weaver, Janeway’s Immunobiology (2016).

14. D. M. Pardoll, The blockade of immune checkpoints in cancer immunotherapy. Nat. Rev. Cancer. 12, 252–64 (2012).

15. M. A. Swartz, S. Hirosue, J. A. Hubbell, Engineering approaches to immunotherapy. Sci Transl Med. 4, 148rv9 (2012).

16. R. F. Sîrbulescu, C. K. Boehm, E. Soon, M. Q. Wilks, I. Ilieş, H. Yuan, B. Maxner, N. Chronos, C. Kaittanis, M. D. Normandin, G. El Fakhri, D. P. Orgill, A. E. Sluder, M. C. Poznansky, Mature B cells accelerate wound healing after acute and chronic diabetic skin lesions. Wound Repair Regen. 25, 774–791 (2017).

17. J. C. Doloff, O. Veiseh, A. J. Vegas, H. H. Tam, S. Farah, M. Ma, J. Li, A. Bader, A. Chiu, A. Sadraei, S. Aresta-Dasilva, M. Griffin, S. Jhunjhunwala, M. Webber, S. Siebert, K. Tang, M. Chen, E. Langan, N. Dholokia, R. Thakrar, M. M. Qi, J. Oberholzer, D. L. Greiner, R. Langer, D. G. Anderson, E. Langman, N. Dholakia, R. Thakrar, M. M. Qi, J. Oberholzer, D. L. Greiner, R. Langer, D. G. Anderson, Colony Stimulating Factor-1 Receptor is a central component of the foriegn body response to biomaterial implants in rodents and non-human primates. Nat. Mater. 16, 671–680 (2017).

18. J. W. Cohen Tervaert, M. J. Colaris, R. R. Van Der Hulst, Silicone breast implants and autoimmune rheumatic diseases: Myth or reality. Curr. Opin. Rheumatol. 29, 348–354 (2017).

19. W. Dolores, R. Christian, N. Harald, P. Hildegunde, W. Georg, Cellular and molecular composition of fibrous capsules formed around silicone breast implants with special focus on local immune reactions. J. Autoimmun. 23, 81–91 (2004).

20. W. E. Katzin, L. J. Feng, M. Abbuhl, M. A. Klein, Phenotype of lymphocytes associated with the inflammatory reaction to silicone gel breast implants. Clin. Diagn. Lab. Immunol. 3, 156–161 (1996).

21. D. W. Oliver, M. Walker, A. Walters, P. Chatrath, B. G. H. Lamberty, Anti-silicone antibodies and silicone containing breast implants. Br. J. Plast. Surg. 53, 410–214 (2000).

22. K. Sadtler, K. Estrellas, B. W. Allen, M. T. Wolf, H. Fan, A. J. Tam, C. H. Patel, B. S. Luber, H. Wang, K. R. Wagner, J. D. Powell, F. Housseau, D. M. Pardoll, J. H. Elisseeff, Developing a pro-regenerative biomaterial scaffold microenvironment requires T helper 2 cells. Science (80-.). 352, 366–370 (2016).

23. L. Chung, D. R. Maestas, A. Lebid, A. Mageau, G. D. Rosson, X. Wu, M. T. Wolf, A. J. Tam, I. Vanderzee, X. Wang, J. I. Andorko, H. Zhang, R. Narain, K. Sadtler, H. Fan, D. Čiháková, C. J. Le Saux, F. Housseau, D. M. Pardoll, J. H. Elisseeff, Interleukin 17 and senescent cells regulate the foreign body response to synthetic material implants in mice and humans. Sci. Transl. Med. 12 (2020), doi:10.1126/scitranslmed.aax3799.

24. P. Ghia, E. Ten Boekel, A. G. Rolink, F. Melchers, B-cell development: A comparison between mouse and man. Immunol. Today. 19, 480–485 (1998).

25. K. Wang, G. Wei, D. Liu, CD19: a biomarker for B cell development, lymphoma diagnosis and therapy. Exp. Hematol. Oncol. 1, 36 (2012).

26. M. Shapiro-Shelef, K. I. Lin, L. J. McHeyzer-Williams, J. Liao, M. G. McHeyzer-Williams, K. Calame, Blimp-1 is required for the formation of immunoglobulin secreting plasma cells and pre-plasma memory B cells. Immunity. 19, 607–620 (2003).

27. S. A. Corfe, C. J. Paige, The many roles of IL-7 in B cell development; Mediator of survival, proliferation and differentiation. Semin. Immunol. 24, 198–208 (2012).

28. T. F. Tedder, A. W. Boyd, A. S. Freedman, S. F. Schlossman, The B cell surface molecule B1 is functionally linked with B cell activation and differentiation. J. Immunol. 135 (2020).

29. T. W. LeBien, T. F. Tedder, B lymphocytes: how they develop and function. Blood. 112, 1570–80 (2008).

30. Y. Feng, N. Seija, J. M. Di Noia, A. Martin, AID in Antibody Diversification: There and Back Again. Trends Immunol. 41, 586–600 (2020).

31. G. Lamson, M. E. Koshland, Changes in J chain and mu chain RNA expression as a function of B cell differentiation. J. Exp. Med. 160, 877–892 (1984).

32. S. F. Badylak, Small intestinal submucosa (SIS): a biomaterial conducive to smart tissue remodeling. Tissue Eng., 179–189 (1993).

33. V. Bronte, J. Pittet, Mikael, The spleen in local and systemic regulation of immunity. Immunity. 39, 806–818 (2013).

34. M. Eikmans, M. C. Roos-Van Groningen, Y. W. J. Sijpkens, J. Ehrchen, J. Roth, H. J. Baelde, I. M. Bajema, J. W. De Fijter, E. De Heer, J. A. Bruijn, Expression of surfactant protein-C, S100A8, S100A9, and B cell markers in renal allografts: Investigation of the prognostic value. J. Am. Soc. Nephrol. 16, 3771–3786 (2005).

35. L. A. N. Crowe, M. McLean, S. M. Kitson, E. G. Melchor, K. Patommel, H. M. Cao, J. H. Reilly, W. J. Leach, B. P. Rooney, S. J. Spencer, M. Mullen, M. Chambers, G. A. C. Murrell, I. B. McInnes, M. Akbar, N. L. Millar, S100A8 & S100A9: Alarmin mediated inflammation in tendinopathy. Sci. Rep. 9, 1–12 (2019).

36. J. Wang, M. Hossain, A. Thanabalasuriar, M. Gunzer, C. Meininger, P. Kubes, Visualizing the function and fate of neutrophils in sterile injury and repair. Science, 358(6359), 111-116. Science (80-.). 358, 111–116 (2017).

37. F. V. Castanheira, P. Kubes, Neutrophils and NETs in modulating acute and chronic inflammation. Blood. 133, 2178–2185 (2019).

38. P. Kruger, M. Saffarzadeh, A. N. R. Weber, N. Rieber, M. Radsak, H. von Bernuth, C. Benarafa, D. Roos, J. Skokowa, D. Hartl, Neutrophils: Between Host Defence, Immune Modulation, and Tissue Injury. PLoS Pathog. 11, 1–22 (2015).

39. A. Beringer, M. Noack, P. Miossec, IL-17 in Chronic Inflammation: From Discovery to Targeting. Trends Mol. Med. 22, 230–241 (2016).

40. L. Huang, D. P. Beiting, N. G. Gebreselassie, L. F. Gagliardo, M. C. Ruyechan, N. A. Lee, J. A. Appleton, Eosinophils and IL-4 support nematode growth coincident with an innate response to tissue injury. PLoS Pathog. 11 (2015), doi:https://doi.org/10.1371/journal.ppat.1005347.

41. Y. S. Goh, N. C. Henderson, J. E. Heredia, A. R. Eagle, J. I. Odegaard, N. Lehwald, K. D. Nguyen, D. Sheppard, L. Mukundan, R. M. Locksley, A. Chawla, Eosinophils secrete IL-4 to facilitate liver regeneration. Proc. Natl. Acad. Sci. 110, 9914–9919 (2013).

42. K. M. M. Vannella, T. A. A. Wynn, K. M. M. Vannella, Macrophages in Tissue Repair, Regeneration, and Fibrosis. Immunity. 44, 450–462 (2016).

43. D. T. Ploeger, N. a Hosper, M. Schipper, J. a Koerts, S. de Rond, R. a Bank, Cell plasticity in wound healing: paracrine factors of M1/ M2 polarized macrophages influence the phenotypical state of dermal fibroblasts. Cell Commun. Signal. 11, 29 (2013).

44. S. Gordon, P. R. Taylor, Monocyte and macrophage heterogeneity. Nat. Rev. Immunol. 5, 953–64 (2005).

45. P. J. Murray, T. A. Wynn, Protective and pathogenic functions of macrophage subsets. Nat. Rev. Immunol. 11, 723–737 (2011).

46. S. F. Badylak, J. E. Valentin, A. K. Ravindra, G. P. McCabe, A. M. Stewart-Akers, Macrophage Phenotype as a Determinant of Biologic Scaffold Remodeling. Tissue Eng. Part A. 14, 1835–1842 (2008).

47. B. Brown, R. Londono, S. Tottey, L. Zhang, K. A. Kukla, M. T. Wolf, K. A. Daly, J. E. Reing, S. F. S. Badylak, Macrophage phenotype as a predictor of constructive remodeling following the implantation of biologically derived surgical mesh materials. Acta Biomater. 8, 978–987 (2012).

48. S. D. Sommerfeld, C. Cherry, R. M. Schwab, L. Chung, D. R. M. Jr, P. Laffont, J. E. Stein, A. Tam, S. Ganguly, F. Housseau, J. M. Taube, D. M. Pardoll, P. Cahan, J. H. Elisseeff, Interleukin-36y – producing macrophages drive IL-17 – mediated fibrosis. Sci. Immunol. 4 (2019), doi:10.1126/sciimmunol.aax4783.

49. M. Rieckmann, M. Delgobo, C. Gaal, L. Büchner, P. Steinau, D. Reshef, C. Gil-Cruz, E. N. ter Horst, M. Kircher, T. Reiter, K. G. Heinze, H. W. M. Niessen, P. A. J. Krijnen, A. M. van der Laan, J. J. Piek, C. Koch, H. J. Wester, C. Lapa, W. R. Bauer, B. Ludewig, N. Friedman, S. Frantz, U. Hofmann, G. C. Ramos, Myocardial infarction triggers cardioprotective antigen-specific T helper cell responses. J. Clin. Invest. 129, 4922–4936 6. (2019).

50. J. T. Walsh, S. Hendrix, F. Boato, I. Smirnov, J. Zheng, J. R. Lukens, S. Gadani, D. Hechler, G. Gölz, K. Rosenberger, T. Kammertöns, J. Vogt, C. Vogelaar, V. Siffrin, A. Radjavi, A. Fernandez-Castaneda, A. Gaultier, R. Gold, T. D. Kanneganti, R. Nitsch, F. Zipp, J. Kipnis, MHCII-independent CD4+ T cells protect injured CNS neurons via IL-4. J. Clin. Invest. 125, 2547 (2015).

51. A. Ulndreaj, Antigona Tzekou, A. J. Mothe, A. M. Siddiqui, R. Dragas, C. Tator, E. E. Torlakovic, M. G. Fehlings, Characterization of the Antibody Response after Cervical Spinal Cord Injury. J. Neurtotrauma. 34, 1209–1226 (2017).

52. R. A. Malt, O. Cope, Antibody peoduction after certain forms of trauma. Surgery. 39, 959–969 (1956).

53. N. Nishio, S. Ito, H. Suzuki, K. Isobe, Antibodies to wounded tissue enhance cutaneous wound healing. Immunology. 128, 369–380 (2009).

54. Y. Iwata, A. Yoshizaki, K. Komura, K. Shimizu, F. Ogawa, T. Hara, E. Muroi, S. Bae, M. Takenaka, T. Yukami, M. Hasegawa, M. Fujimoto, Y. Tomita, T. F. Tedder, S. Sato, CD19, a response regulator of B lymphocytes, regulates wound healing through hyaluronan-induced TLR4 signaling. Am. J. Pathol. 175, 649–660 (2009).

55. K. Yanaba, A. Yoshizaki, Y. Asano, T. Kadono, T. F. Tedder, S. Sato, IL-10-producing regulatory B10 cells inhibit intestinal injury in a mouse model. Am. J. Pathol. 178, 735–743 (2011).

56. G. Sun, Y. Wang, Y. Ti, J. Wang, J. Zhao, H. Qian, Regulatory B cell is critical in bone union process through suppressing proinflammatory cytokines and stimulating Foxp3 in Treg cells. Clin. Exp. Pharmacol. Physiol. 44, 455–462 (2017).

57. S. Yang, W. Ding, H. Feng, H. Gong, D. Zhu, B. Chen, J. Chen, Loss of B cell regulatory function is associated with delayed healing in patients with tibia fracture. APMIS. 123, 975–985 (2015).

58. R. M. Goldblum, A. A. O’Donell, D. Pyron, R. P. Pelley, J. P. Heggers, Antibodies to silicone elastomers and reactions to ventriculoperitoneal shunts. Lancet. 340, 510–513 (1992).

59. L. E. Wolf, M. Lappé, R. D. Peterson, E. G. Ezrailson, Faseb J, in press, doi:https://doi.org/10.1096/fasebj.7.13.8405812.

60. E. Karlson, S. Hankinson, M. Liang, J. Sanchez-Guerrero, G. Colditz, B. Rozenau, F. Speizer, P. Schur, Association of silicone breast implants with immunologic abnormalities: a prospective study. Am J Med, 1999;106:11–9. Am. J. Med. 106, 11–19 (1999).

61. K. L. White, P. C. Klykken, The Non-Specific Binding of Immunoglobulins to Silicone Implant Materials: The Lack of a Detectable Silicone Specific Antibody. Immunol. Invest. 27, 221–235 (1998).

62. R. Herdman, V. Ernster, S. Bonurant, Safety of Silicone Breast Implants (2000).

63. J. E. Babensee, M. V. Sefton, Viability of HEMA-MMA Microencapsulated Model Hepatoma Cells in Rats and the Host Response. Tissue Eng. 6 (2004), doi:http://doi.org/10.1089/107632700320784.

64. D. M. Lindell, A. A. Berlin, M. A. Schaller, N. W. Lukacs, B cell antigen presentation promotes Th2 responses and immunopathology during chronic allergic lung disease. PLoS One. 3 (2008), doi:10.1371/journal.pone.0003129.

65. P. J. Linton, B. Bautista, E. Biederman, E. S. Bradley, J. Harbertson, R. M. Kondrack, R. C. Padrick, L. M. Bradley, Costimulation via OX40L expressed by B cells is sufficient to determine the extent of primary CD4 cell expansion and TH2 cytokine secretion in vivo. J. Exp. Med. 197, 875–883 (2003).

66. S. Takemura, P. A. Klimiuk, A. Braun, J. J. Goronzy, C. M. Weyand, T Cell Activation in Rheumatoid Synovium Is B Cell Dependent. J. Immunol. 167, 4710–4718 (2001).

67. X. Chen, P. E. Jensen, The role of B lymphocytes as antigen-presenting cells. Arch. Immunol. Ther. Exp. (Warsz). 56, 77–83 (2008).

68. B. M. Sicari, V. Agrawal, B. F. Siu, C. J. Medberry, C. L. Dearth, N. J. Turner, S. F. Badylak, A murine model of volumetric muscle loss and a regenerative medicine approach for tissue replacement. Tissue Eng. - Part A. 18, 1941–1948 (2012).

69. I. Tirosh, B. Izar, S. M. Prakadan, M. H. Wadsworth, D. Treacy, J. J. Trombetta, C. Rodman, C. Lian, G. Murphy, M. Fallahi-Sichani, Dissecting the multicellular ecosystem of metastatic melanoma by single-cell RNA-seq. Science (80-.). 352, 189–196 (2016).

